# Molecular and histologic outcomes following spinal cord injury in spiny mice, *Acomys cahirinus*

**DOI:** 10.1101/793133

**Authors:** K.A. Streeter, M.D. Sunshine, J.O. Brant, M. A.G.W. Sandoval, M. Maden, D.D. Fuller

**Affiliations:** Department of Physical Therapy, University of Florida, Gainesville, Florida 32601; McKnight Brain Institute, University of Florida, Gainesville, Florida 32601; Center for Respiratory Research and Rehabilitation, University of Florida, Gainesville, FL 32610; Department of Biology, University of Florida Gainesville FL 32610

## Abstract

The spiny mouse (*Acomys cahirinus)* appears to be unique among mammals by showing little scarring or fibrosis after skin or muscle injury, but the *Acomys* response to spinal cord injury (SCI) is unknown. We tested the hypothesis that *Acomys* would have molecular and immunohistochemical evidence of reduced spinal inflammation and fibrosis following SCI as compared to C57BL/6 mice (*Mus*), which similar to all mammals studied to date exhibits spinal scarring following SCI. Initial experiments used two pathway-focused RT-PCR gene arrays (“wound healing” and “neurogenesis”) to evaluate tissue samples from the C2-C6 spinal cord 3-days after a C3/C4 hemi-crush injury (C3Hc). Based on the gene array, specific genes were selected for RT-qPCR evaluation using species-specific primers. The results supported our hypothesis by showing increased inflammation and fibrosis related gene expression (*Serpine 1, Plau, Timp1)* in *Mus* as compared to *Acomys* (P<0.05). RT-qPCR also showed enhanced stem cell and axonal guidance related gene expression *(Bmp2, GDNF, Shh)* in *Acomys* compared to *Mus* (P<0.05). Immunohistochemical evaluation of the spinal lesion at 4-wks post-injury indicated reduced collagen IV immunostaining in *Acomys* (P<0.05). Glial fibrillary acidic protein (GFAP) and ionized calcium binding adaptor molecule 1(IBA1) immunostaining indicated morphological differences in the appearance of astrocytes and macrophages/microglia in *Acomys*. Collectively, the molecular and histologic results support the hypothesis that *Acomys* has reduced spinal inflammation and fibrosis following SCI. We suggest that *Acomys* may be a useful comparative model to study adaptive responses to SCI.

**Highlights:**

- Spiny mice (*Acomys cahirinus)* and C57BL/6 (*Mus*) were studied after spinal injury
- RT-PCR gene arrays suggested different molecular response in *Acomys*
- RTq-PCR with species-specific primers showed increased neurogenesis-related signaling
- Histology indicates reduced scarring and fibrosis in *Acomys*
- *Acomys* may be a useful comparative model to study SCI

## Introduction

A fundamental goal of regenerative medicine is to develop therapies to reduce inflammation, fibrosis, scarring, or other detrimental outcomes in damaged or diseased mammalian tissues. Spinal cord injury (SCI) is one condition in which inflammation, scarring, and limited regeneration has severe functional consequences. Immediately after SCI an inflammatory response is induced involving activation of resident immune cells (e.g. microglia), and recruitment of monocyte/macrophages and neutrophils to the site of the injury (Carlson, Parrish, Springer, Doty, & Dossett, 1998; Hawthorne & Popovich, 2011). At later time points, a glial/fibrotic scar forms and provides a barrier that prevents axonal regeneration (Tran, Warren, & Silver, 2018). Current regenerative therapies in SCI include delivery of exogenous stem cells (Pereira, Marote, Salgado, & Silva, 2019) and/or the incorporation of biomolecular scaffolds into the damaged spinal cord (Kim, Park, & Choi, 2014). However, regenerative failure remains a consistent feature of SCI in animal models and humans.

One contributing factor to the lack of progress in identifying therapeutic targets has been the absence of a relevant model system, namely an adult mammal with high regenerative capacity of spinal tissue. In this regard, we recently discovered that the adult spiny mouse (*Acomys cahirinus*) can regenerate a remarkable range of tissues following injury including dermis, hair follicles, sebaceous glands, smooth muscle, skeletal muscle, adipose cells and cartilage (Brant, Yoon, Polvadore, Barbazuk, & Maden, 2016; Seifert et al., 2012). *Acomys* does not undergo fibrosis in response to damage in any organ examined thus far, and presents a unique opportunity to interrogate the mechanisms involved in mammalian tissue regeneration (Pinheiro, Prata, Araújo, & Tiscornia, 2018). A mammalian model of improved regenerative capacity after SCI would provide an opportunity for comprehensive examination of the mechanisms contributing to this response. Towards this goal, we evaluated the acute molecular and long-term immunohistochemical impact of SCI in *Acomys*. This response was compared to *Mus*, which displays signatures of the “typical” mammalian response to spinal injury (i.e. inflammation, scarring, limited regeneration). The overall hypothesis was that the injured spinal cord in adult *Acomys* would have a molecular and immunohistochemical signature consistent with reduced fibrosis and improved regenerative capacity as compared to *Mus*.

## Materials and methods

### Animals

All experiments were conducted with adult, male C57/BL6 (n=16; 29.8 ± 2.8g) mice (*Mus*; Jackson Laboratory) and male, spiny mice (n=15; 45.1 ± 6.4g) (*Acomys cahirinus*; in-house colony at the University of Florida). Descriptions of skin and muscle wound healing using spiny mice from this colony have been published (Brant, Lopez, Baker, Barbazuk, & Maden, 2015; Brant et al., 2016; Maden et al., 2018). Mice were housed in a controlled environment (12h light/dark cycles) with food and water *ad libitum*. All experimental protocols were approved by the Institutional Animal Care and Use Committee at the University of Florida.

### Spinal Cord Injury

Anesthesia was induced by placing mice in a chamber flushed with 3% isoflurane mixed with 100% O_2_. Anesthetized mice were transferred to a heated surgical station and core body temperature was maintained at 37.5 ± 0.5°C (model 700 TC-1000, CWE). During surgery, anesthesia was maintained by having the mice breathe 1.5-2% isoflurane through a nose cone with 100% O_2_. The surgical area was cleaned with three, alternating rounds of betadine surgical scrub followed by 70% ethanol. A dorsal incision was then made over the spinal midline from the base of the skull to the fifth cervical segment. A C3 laminectomy was performed to expose the spinal cord, and a C3 lateral, dorsal crush injury was induced similar to previous report (Hilton et al., 2013). A sterilized pair of fine tipped Dumont #5 forceps was marked 1mm from the tip with a black sharpie marker. To induce the injury, one prong of the forceps was inserted at the spinal midline at a depth of 1mm (identified by the black sharpie mark) while the other prong remained outside the lateral edge of the spinal cord. The forceps were held closed for 15 seconds. The forceps were removed, reinserted for a second time and closed again for 15 seconds. Following the injury, the overlying muscles were sutured with 4-0 Vicryl suture and the skin was closed with sterile wound clips. Mice received an analgesic, buprenorphine (0.03 mg/kg, s.q.) and nonsteroidal anti-inflammatory medication meloxicam (2.0 mg/kg, s.q.) for the initial 48 hours post-injury. Mice also received Lactated Ringer’s solution (2 ml/day, s.q.) and oral Nutri-cal supplements (0.5-1ml, Webster Veterinary, MA, USA) until adequate volitional eating and drinking resumed. Manual bladder expression was performed twice daily until voluntary micturition was apparent. The bladder size was estimated by palpating the lower abdomen using two fingers (thumb and index finger) and was then expressed by gently applying pressure until the bladder was completely emptied. Body temperature was assessed daily for two weeks post-SCI using a digital mouse rectal thermometer (model 700 TC-1000, CWE) and Vaseline.

### Gene Arrays

Three days post-SCI (*Mus*: n=6, *Acomys*: n=5) or age matched controls not receiving an injury (*Mus*: n=4, *Acomys*: n=5), mice were anesthetized by breathing 3% isoflurane mixed into 100% O_2_. The mice then received an i.p. injection of beuthanasia solution (150mg/kg). The depth of anesthesia was confirmed via lack of foot withdrawal to toe pinch, and the cervical spinal cord from C2 to C6 was quickly extracted, placed in an ice cold RNA free dish, cut sagittally to isolate the ipsilateral (same side as injury) spinal cord and immediately placed into 1.5mL of RNALater (Qiagen Cat. 76104). Spinal cord samples were stored at 4°C for 24 hours and then maintained at −80°C. Tissues were subsequently thawed at 4°C, washed in RNase-free water and homogenized using a rotor stator type tissue homogenizer (ProScientific Bio-Gen PRO200 Homogenizer; Multi-Gen 7XL Generator Probes) in RLT Buffer and processed using the Rneasy Plus Mini Kit (Qiagen Cat. 74134). RNA quality was assessed using an Agilent 2200 TapeStation (Agilent, Andover, MA). All samples had a RIN score > 7.0. For the wound healing pathway-focused RT-PCR array (Qiagen PAMM-121Z) and the neurogenesis pathway-focused RT-PCR array (Qiagen PAMM-404Z) cDNA was generated using the RT_2_ PreAMP cDNA Synthesis Kit (Qiagen 330451), using array specific primers (Qiagen 330241) followed by the RT^2^ First Strand Kit (Qiagen 33041). Arrays were run using RT^2^ SYBR Green qPCR Mastermix (Qiagen 330502). For RT-qPCR with species-specific primers cDNA was generated SuperScript™ IV VILO Reverse Transcriptase (Invitrogen 11756050) following the manufacturer’s protocol. Real-Time PCR was performed using Sso-Fast™ EvaGreen® Supermix (Bio-Rad 172-5200) on a Bio-Rad C1000 Touch™ Thermal Cycler. The fold change in gene expression was calculated using the ΔΔCt relative expression method (Livak & Schmittgen, 2001) using *Gapdh* as the reference gene. Sequence of species-specific PCR primers can be found in Supplemental Table 1. All reactions were run with an annealing temperature of 60°C.

**Table 1:** Antibody name, source, catalog number, concentration, immunogen, and research resource identifiers (RRID) for each primary antibody used in the study.

| Antibody | Source | Catalog # | Concentration | Immunogen | RRID |
| --- | --- | --- | --- | --- | --- |
| Collagen IV | Abcam | ab6586 | 1:250 | Full length native protein (purified) corresponding to Collagen IV from human and bovine placenta | AB_305584 |
| MBP | Encor Biotechnology | CPCA-MBP | 1:500 | Purified myelin basic protein isolated from bovine brain | AB_2572352 |
| Iba1 | Wako Chemicals | 019-19741 | 1:300 | Synthetic peptide (C-terminal of Iba1) | AB_839504 |
| NeuN | Encor Biotechnology | MCA-1B7 | 1:1000 | N-terminal 99 amino acids of human FOX3 expressed in and purified from E. coli | AB_2572267 |
| GFAP | Encor Biotechnology | CPCA-GFAP | 1:1000 | Recombinant full length human GFAP isotype 1 expressed in and purified from E. coli | AB_2109953 |

### Antibody Selection

Antibody name, source, catalog number, concentration, immunogen, and research resource identifiers (RRID) for each primary antibody used in the study are provided in Table 1. The polyclonal, Collagen IV antibody (Abcam, Cat#ab6586; RRID: AB_305584; sequence available here: https://www.uniprot.org/uniprot/P02462) was validated by the manufacturer and binds to native collagen epitopes composed of multiple subunit strands and has negligible cross-reactivity with Type I, II, III, V or VI collagens. Specifically, Collagen IV has been shown to accumulate within the scar following SCI (Liesi & Kauppila, 2002). This antibody detects a single band ∼250kDa on western blots in baby hamster kidney fibroblasts (Abcam, 2012), and has an expression pattern following SCI in mouse (Vangansewinkel et al., 2019) and rat (Tuinstra et al., 2014) similar to that reported herein.

The polyclonal, Myelin Basic Protein (MBP) antibody (EnCor Biotechnology, Cat#CPCA-MBP; RRID: AB_2572352; sequence available here: http://encorbio.com/Alignments/MBP%20isoforms.pdf binds to all four gene products from the single mammalian MBP gene which is observed by four bands between ∼14-21.5 kDa in western blots in rodent spinal tissue (EnCor Biotechnology). The Iba-1 antibody (Wako Chemicals, Cat#019-19741; RRID: AB_839504) recognizes a calcium-binding protein with a molecular weight of 17 kDa specifically expressed in macrophage/microglia (Ito et al., 1998) and has been validated for immunohistochemistry in rodent, human, dog, cat, pig, marmoset, and zebrafish (Ahn et al., 2012; Fantin et al., 2010; Gaige et al., 2013; Ide, Uchida, Tamura, & Nakayama, 2010; Rodriguez-Callejas, Fuchs, & Perez-Cruz, 2016; S. M. Turner et al., 2016). The Neu-N antibody (EnCor Biotechnology, Cat#MCA-1B7; RRID: AB_2572267; sequence available here: https://www.uniprot.org/uniprot/A6NFN3) is a reliable neuronal marker, binding to neurons in all vertebrates (Mullen, Buck, & Smith, 1992). We have previously used the polyclonal, Glial Fibrillary Acidic Protein (GFAP) antibody (EnCor Biotechnology, Cat#CPCA-GFAP; RRID: AB_2109953; sequence available here: https://www.uniprot.org/uniprot/P14136) to stain reactive astrocytes in mouse brain and spinal tissue during CNS disease (S. M. Turner et al., 2016; Turner, Falk, Byrne, & Fuller, 2016). Similar to other reports (Bovolenta, Wandosell, & Nieto-Sampedro, 1992; Cregg et al., 2014), GFAP is a major constituent of the astro-glial scar in most mammals which forms following SCI. Western blot analysis of whole brain lysates indicates GFAP antibody shows a single, strong band at ∼50kDa.

### Immunohistochemistry

A separate cohort of mice that recovered for four weeks following SCI mice (*Mus*: n=6, *Acomys*: n=5) were anesthetized by breathing 3% isoflurane mixed into 100% O_2_ followed by an i.p. injection of urethane solution (1.0-1.6g/kg in distilled water). After the depth of anesthesia was verified, mice were transcardially perfused with ice cold saline (1ml per gram body weight) followed by 4% paraformaldehyde (PFA; 1ml/g) in 1X Dulbecco’s phosphate buffered saline (DPBS, Mediatech, Inc., Cat#21-030-CV). The cervical spinal cord (C2-C6) was harvested, post-fixed in 4% PFA overnight at 4°C and placed in 70% ethanol at room temperature for at least 48 hrs. The spinal cord was then paraffin embedded and stored at room temperature. Paraffin embedded spinal cords were cut longitudinally at 7 µm on a Leica Biosystems microtome and mounted on glass slides (Fisher, Superfrost Plus). Alternate slides were then used for immunostaining, and this enabled each stain to be evaluated at 35μm increments (i.e., every 5^th^ slide was stained).

Prior to immunochemistry, tissue sections were deparaffinized and rehydrated through xylenes followed by a graded series of ethanol exposures (100%, 95% and 70%). An antigen retrieval procedure was done using 0.1M Citrate solution, pH 6.0 for 25 minutes (collagen IV/MBP staining; Abcam) or Trilogy reagent (IBA1/GFAP/NeuN staining; Cell Marque) at 95 °C for 15 min. Slides were blocked in 2% normal horse serum (Vector Labs) in 1X TBS with 0.2% Triton-X for 1hr at room temperature. Spinal sections were then incubated for two days at 4°C in the primary antibody solution (Table1): rabbit anti-collagen IV (1:250; RRID: AB_305584), chicken anti-MBP (1:500; RRID: AB_2572352), rabbit anti-Iba-1 (1:300; RRID: AB_839504), mouse anti-NeuN (1:1,000; RRID: AB_2572267), chicken anti-GFAP (1:1,000; RRID: AB_2109953), in antibody diluent (ThermoFisher Scientific, #003118). Immunoreactivity was detected using Alexa Fluor® secondary antibodies (all 1:500; ThermoFisher Scientific) donkey anti-mouse 488 (Cat#A-21202), donkey anti-rabbit 594 (Cat#A-21207), goat anti-chicken 488 (Cat#A-11039), and goat anti-chicken 594 (Cat#A-11042) in 2% normal horse serum in 1X TBS with 0.2% Triton-X for 1hr at room temperature. Positive control tissues and negative control tissue concentration matched Ig controls were included with each immunoassay. Negative controls resulted in no staining. Slides were coverslipped with VectaShield antifade mounting medium containing DAPI (Vector Labs, Cat#H-1200). Slides were air-dried and stored at 4°C. All fluorescent images were captured with a BZ-X series all-in-one fluorescence microscope (Keyence Corporation).

### Data and Statistical Analyses

Gene expression 3 days post-SCI was compared to spinal intact controls within each species and genes with a fold change >1.5 are provided in Supplemental Table 2. Individual one tailed t-tests were used for analysis of species-specific primer results.

For immunohistochemistry analysis, images were captured at 10X, stitched together using Adobe Photoshop software and analyzed using MATLAB (The MathWorks R2015a). Briefly, MATLAB code was written to 1) detect the total number of pixels in the section, 2) remove the background (determined empirically based on a blinded subset of images), and 3) calculate the average fluorescent intensity and number of the remaining pixels. Quantification of all immunohistochemistry was performed on every third stained spinal cord section (*Mus*: n=11 ± 0.2; *Acomys*: n=14 ± 0.6 sections). This represented 105μm increments from the dorsal surface of the spinal cord. For GFAP and IBA1 staining, quantification was performed on the entire ipsilateral spinal cord. Collagen IV and MBP staining was quantified at the site of lesion. For quantification, an area of interest that was 4.7% of the tissue section area was positioned to encompass all the collagen IV staining. This value was empirically determined by assessing the spinal lesion from all animals based on the distribution of collagen IV staining. Immunostaining for GFAP and IBA1 was normalized to the size of the ipsilateral (to SCI) tissue section (% positive). To assess immunostaining according to the location of the section within the dorsal-ventral neuroaxis, the total number of imaged sections for each animal was divided into three equal groups representing ventral, middle, and dorsal spinal regions. The average number of sections in each region was similar: ventral: *Mus*: n=3.6 ± 0.2; *Acomys*: n=4.6 ± 0.2 sections, middle: *Mus*: n=3.4 ± 0.2; *Acomys*: n=4.6 ± 0.2 sections, and dorsal: *Mus*: n=3.6 ± 0.2; *Acomys*: n=4.6 ± 0.2 sections. Quantitation of the staining intensity for various makers of the SCI response (e.g., collagen IV, MBP, GFAP, IBA1) was done by evaluating both the raw signal intensity captured via fluorescence microscopy (arbitrary units) and also by normalization relative to the total amount of positive staining that could detected. Statistical analyses were performed in GraphPad Prism 8 (GraphPad Software, Inc). Means are presented with standard error.

### Data availability statement

The data reported herein are available from the corresponding author (DDF) upon reasonable request.

## Results

### Behavioral observations

Several relevant observations were made during the routine care of the animals following SCI (Figure 1). First, *Acomys* resumed spontaneous bladder voiding earlier than *Mus* (Figure 1A). *Acomys* regained the ability to void the bladder by two days post-injury whereas *Mus* required manual bladder expression for more than two weeks. Second, both species showed a transient weight loss after the injury, but with a different time course as illustrated in Figure 1B. *Mus* showed an immediate drop in body mass, whereas *Acomys* did not drop body mass until day three. Prior to injury, body temperature was different between *Mus* (n=6; 37.6±0.1°) and *Acomys* (n=7; 35.7±0.2°, P<0.0001). Differences were also noted in rectal temperature over the first few days following the injury. *Acomys* showed an increase of approximately 1°C during days 1-7 post-injury whereas *Mus* slightly dropped temperature (Figure 1C).

**Figure 1:**
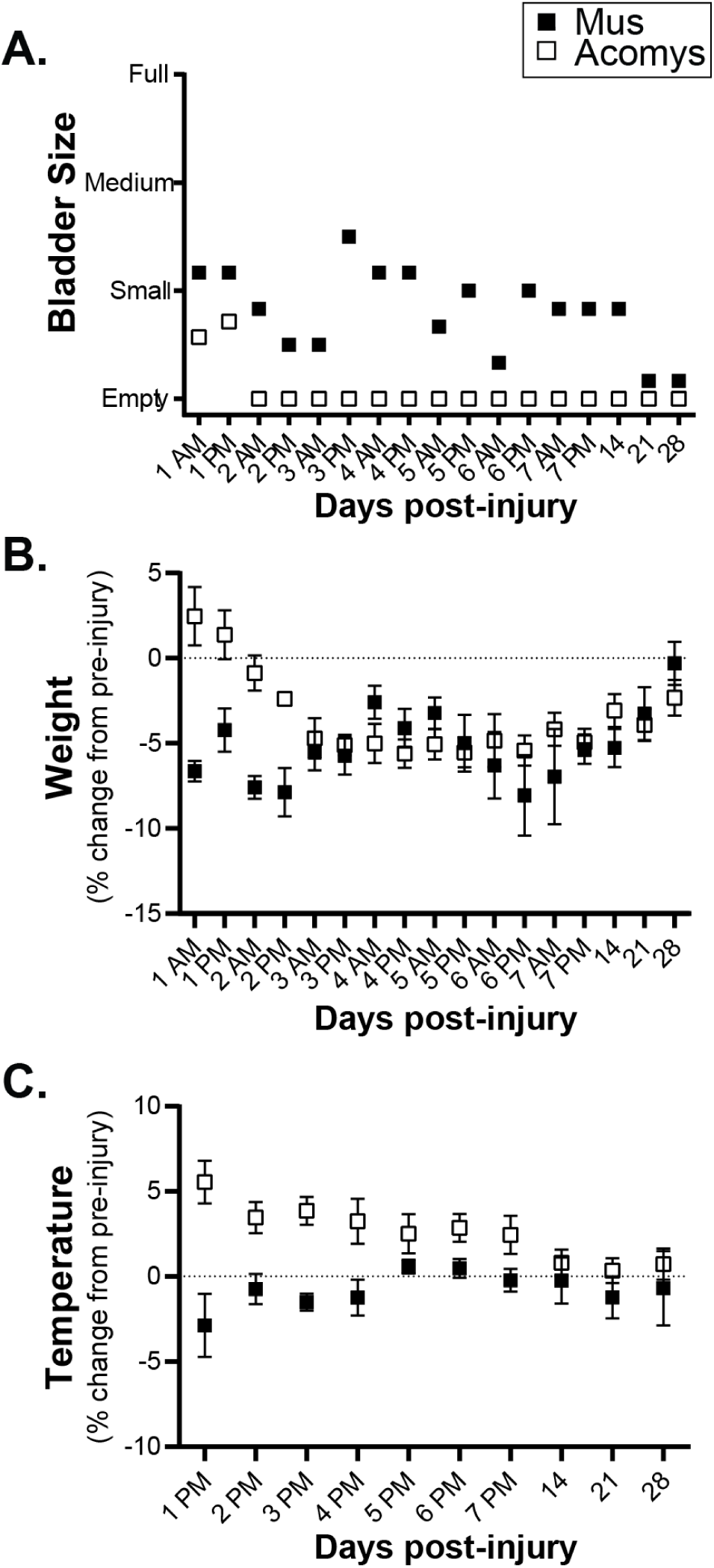
Behavioral observations for 28 days post-SCI. **A**. Bladder size in *Mus* (shaded squares) and *Acomys* (open squares) measured by palpating the lower abdomen. **B**. Weight (as a percent change from pre-injury values) assessed twice a day for the first 7 days and at 2, 3, and 4 weeks post-injury. **C**. Rectal temperature (as a percent change from pre-injury values) assessed once a day for the first 7 days and at 2, 3, and 4 weeks post-injury. Data shown are means ± standard error.

### RT-PCR arrays

For an initial comparison of the molecular genetic response to SCI in *Mus* vs. *Acomys*, we examined the expression profile of the ipsilesional cervical spinal cord (C2-C6) at three days post-SCI (relative to non-injured controls) using two different pathway-focused RT-PCR arrays. These initial experiments were intended to identify genes of interest and individual genes were subsequently validated with RT-qPCR quantitation using both *Mus*-specific and *Acomys*-specific primers (next section). The first gene array targeted 84 genes involved in wound healing (e.g., inflammation, granulation, tissue remodeling) and the second array evaluated 84 genes associated with neurogenesis (e.g., proliferation, differentiation, motility, migration) and stem cell differentiation.

The results of the wound healing array and neurogenesis array are presented in Supplemental Table 2. The wound healing array indicated *Mus* had a stronger wounding response compared to *Acomys*. Specifically, *Mus* had 16 upregulated genes (>1.5 fold) and 21 downregulated compared to expression in the non-injured spinal cord; and *Acomys* had three upregulated and 24 downregulated genes (Supplemental Table 2a). The upregulated genes in *Mus* can be associated into four groups: 1) pro-inflammatory cytokines involved in the immune response to wounding (*Il6, Cxcl3, Ccl12, Ccl7, Tnf*, and *Il1b*); 2) ECM and its remodeling (*Timp1, Itga5, Plaur, Mmp1a, Col5a3*, and *Plau*); 3) Tgfβ1 signaling and fibrosis (*Serpine1, Tagln* and *Tgfb1*); and 4) a growth factor (*Hgf*). In contrast, upregulated genes in *Acomys* consisted of one inflammatory response gene (*Tnf*) and two genes associated with the ECM and cell surface (*Col 5a1* and *Cadherin1*). The majority of gene changes in *Acomys* were in the negative direction including extracellular matrix constituents (*Vtn* and *Clo4a3*), a vitronectin receptor (*Itgav)*, inflammatory cytokines (*Cxcl3 and Il2*) and receptor (*Il6st)*, and two integrins (*Itgb5* and *Itgb6)*.

In stark contrast to the wounding arrays, the majority of the upregulated genes in neurogenesis arrays were observed in *Acomys*. Specifically, there were 22 upregulated genes and three downregulated in *Acomys*, whereas in *Mus* there were seven upregulated genes and 42 downregulated (Supplemental Table 2b). The upregulated genes in *Mus* were *Tgfb1*, two growth factors (*Gdnf* and *Fgf2*), a transcriptional activator of growth factors (*Stat3*), a neuregulin receptor (*Erbb2*), and two genes involved in the regulation of actin cytoskeleton and cell motility (*S100a6* and *Flna*). Four of these seven were also upregulated in *Acomys*. The 22 upregulated genes in the *Acomys* spinal cord included: 1) growth factors/transcriptional activator of growth factors (*Bdnf, Gdnf, Ptn, Stat3); 2) WNT* signaling molecules *(Dvl3, Ndp, Shh*); 3) neural stem cell genes/transcription factors (*Notch1, Notch2, Creb1, Sox2, Ascl1, Heyl* and *Mef2c*); 4) genes involved in axonal guidance (*Ntn1, Robo1*, and *Efnb1*); 5) Tgfβ1 signaling molecules (*Tgfb1, Bmp2);* and 6) a cell death gene (*Bcl2*).

### RT-qPCR of select genes using species-specific primers

Based on the results of the gene arrays, we selected and verified the expression of four genes associated with wound healing and fibrosis in *Mus* and *Acomys* following SCI: *Serpine1, Plau, Timp1*, and *Itgb5* (Figure 2). The levels of mRNA for each gene are expressed relative to non-injured spinal tissue of each species. *Serpine 1*, a major physiological regulator of the plasmin-based cascade involved in fibrotic disorders and pro-inflammatory molecule (Gupta, Xu, Castellino, & Ploplis, 2016) was upregulated post-SCI to a greater extent in *Mus* than *Acomys* (t(4)=6.441, P=0.0015). *Plau*, another plasminogen activator involved in the fibrotic pathway (He, Tsou, Khanna, & Sawalha, 2018) was also upregulated in *Mus* (t(4)=2.516, P=0.0328; Figure 2b). Analysis of *Timp1*, an extracellular matrix enzyme which functions to inhibit the action of the matrix metalloproteases was highly upregulated in *Mus* (t(4)=27.75, P<0.0001; Figure 2c.). *Itgb5*, an integrin that interacts with the matrix and intracellular compartments was similar between *Mus* and *Acomys*, which was supported by statistical analysis (t(4)=1.405, P=0.1163; Figure 2d).

**Figure 2:**
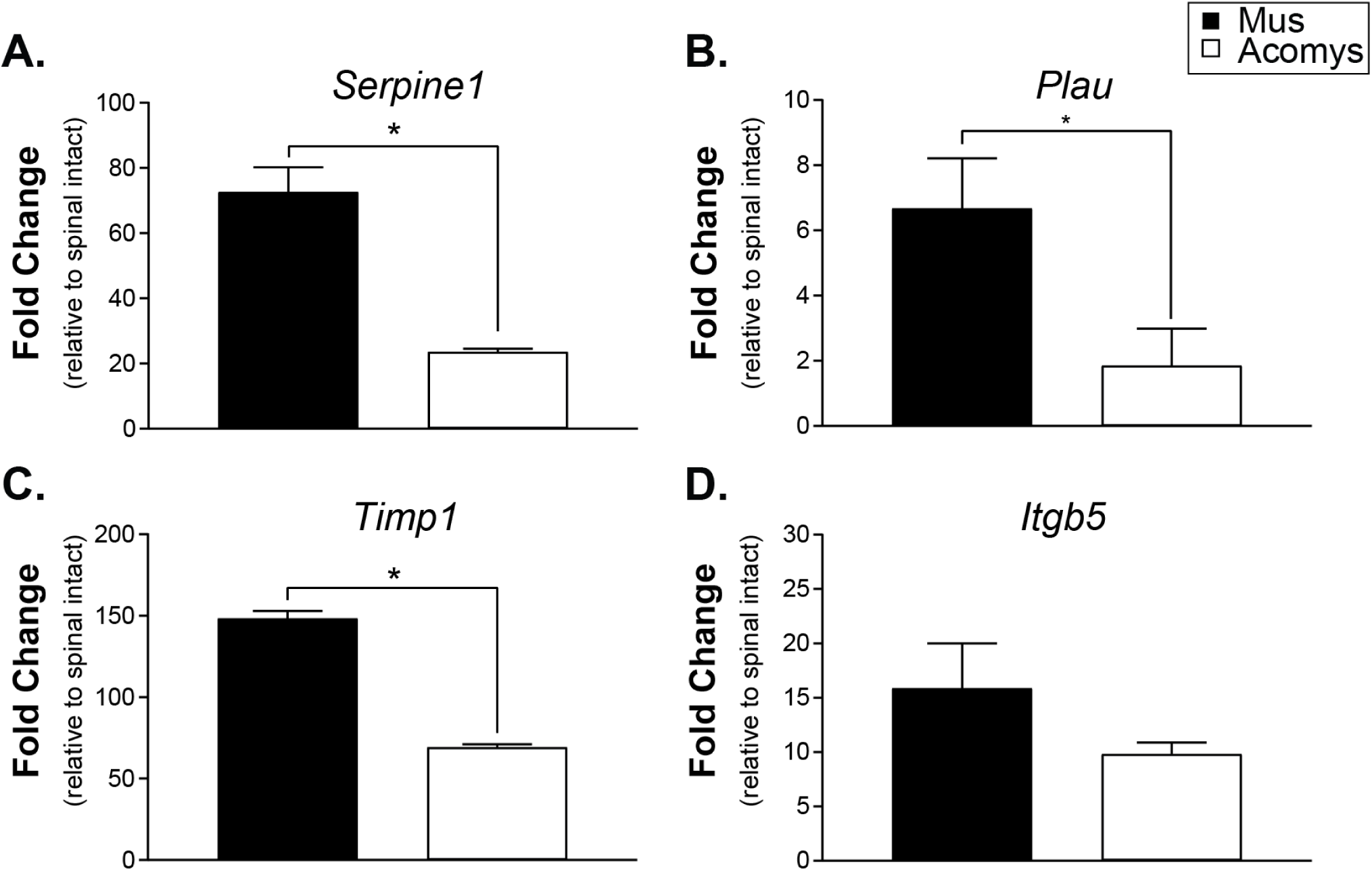
Gene expression levels at 3 days post-SCI expressed as fold change compared to spinal intact controls determined by RT-qPCR using species-specific primers. **A**. *Serpine1* (plasminogen activator inhibitor 1). **B**. *Plau* (plasminogen activator urokinase). **C**. *Timp1* (tissue inhibitor of metalloprotease 1). **D**. *Itgb5* (integrin β5). Data shown are means ± standard error. * = P < 0.05.

We also selected eight growth factor, neural stem cell, and signaling molecules genes identified by the neurogenesis array and verified their expression in both *Mus* and *Acomys* following SCI: *Bmp2, GDNF, Shh, Tgfβ1, Stat3, Notch1, Notch2, and Sox2* (Figure 3). *Bmp2* and *GDNF*, which regulate cellular differentiation (Xiao, Du, Wu, & Yip, 2010) and growth (Rosich, Hanna, Ibrahim, Hellenbrand, & Hanna, 2017) were upregulated in *Acomys* compared to *Mus* ((t(4)=4.936, P=0.0039; Figure 3a) and (t(4)=4.458, P=0.0056; Figure 3b)). *Shh*, a morphogen that aids in progenitor proliferation (Bambakidis, Wang, Franic, & Miller, 2003) and myelination (Thomas & Shea, 2013) was also upregulated in *Acomys* (t(4)=3.883, P=0.0089; Figure 3c). As observed in both gene arrays, levels of *Tgfβ1* were similar between *Mus* and *Acomys* (t(4)=0.7728, P=0.2414; Figure 3d). Although a trend for upregulation, there was no difference in the transcription factor *Stat3* (t(4)=1.696, P=0.0826; Figure 3e). *Notch* signaling, known to regulate neural progenitor cell fate (Cardozo, Mysiak, Becker, & Becker, 2017) was similar between *Mus* and *Acomys* (*Notch1:* (t(4)=0.8279, P=0.2271; Figure 3f); *Notch2:* (t(4)=0.2466, P=0.4087; Figure 3g)). *Sox2*, a transcription factor related to neuronal differentiation was not different between species (t(4)=1.373, P=0.1208; Figure 3g).

**Figure 3:**
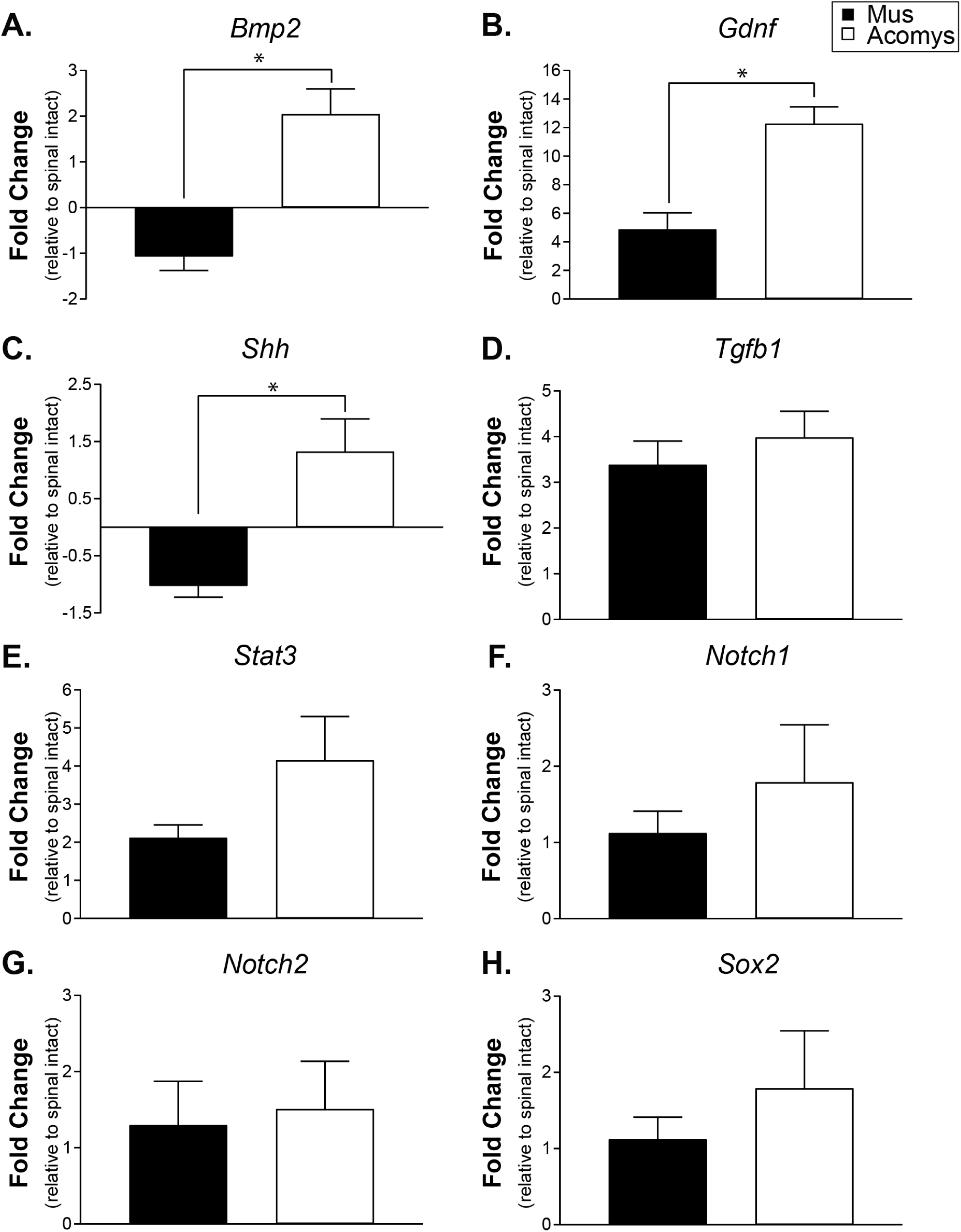
Gene expression levels at 3 days post-SCI expressed as fold change compared to spinal intact controls determined by RT-qPCR using species-specific primers. **A**. *Bmp2* (bone morphogenetic protein 2). **B**. *Gdnf* (glial derived neurotrophic factor). **C**. *Shh* (sonic hedgehog). **D**. *Tgfb1* (transforming growth factor β1). **E**. Stat3 (signal transducer and activator of transcription 3). **F**. *Notch1* (Notch homolog 1, translocation-associated). **G**. *Notch2* (Notch homolog 2, translocation-associated). **H**. *Sox2* (SRY (sex determining region Y)-box 2). Data shown are means ± standard error. * = P < 0.05.

### Spinal cord immunohistochemistry

Representative photomicrographs of lesioned spinal cords stained for collagen IV and MBP are provided in Figure 4A-F. Example “heat maps” which illustrate the density of these markers are shown in Figure 4G-H. Evaluation of the immunohistochemical staining indicated a reduction in the density of collagen IV staining within the spinal lesion in *Acomys*, and this conclusion is supported by the statistical analyses. Evaluation of signal intensity (a.u.) produced low P-values for both species (*Mus vs. Acomys*, F(1,27) = 17.11, P=0.0003) and location (i.e., dorsal, middle, ventral, F(2,27) = 22.87, P<0.0001). Evaluation of normalized staining (% of the lesion area that stained positive for collagen IV) also produced low P-values: species: F(1,27)=23.37, P<0.0001; location: F(2,27)=22.70, P<0.0001. Post-hoc evaluation of the normalized data confirms a difference between *Acomys* and *Mus* in the dorsal (P=0.001), middle (P=0.025) and ventral (P=0.028) spinal regions (Figure 4J).

**Figure 4:**
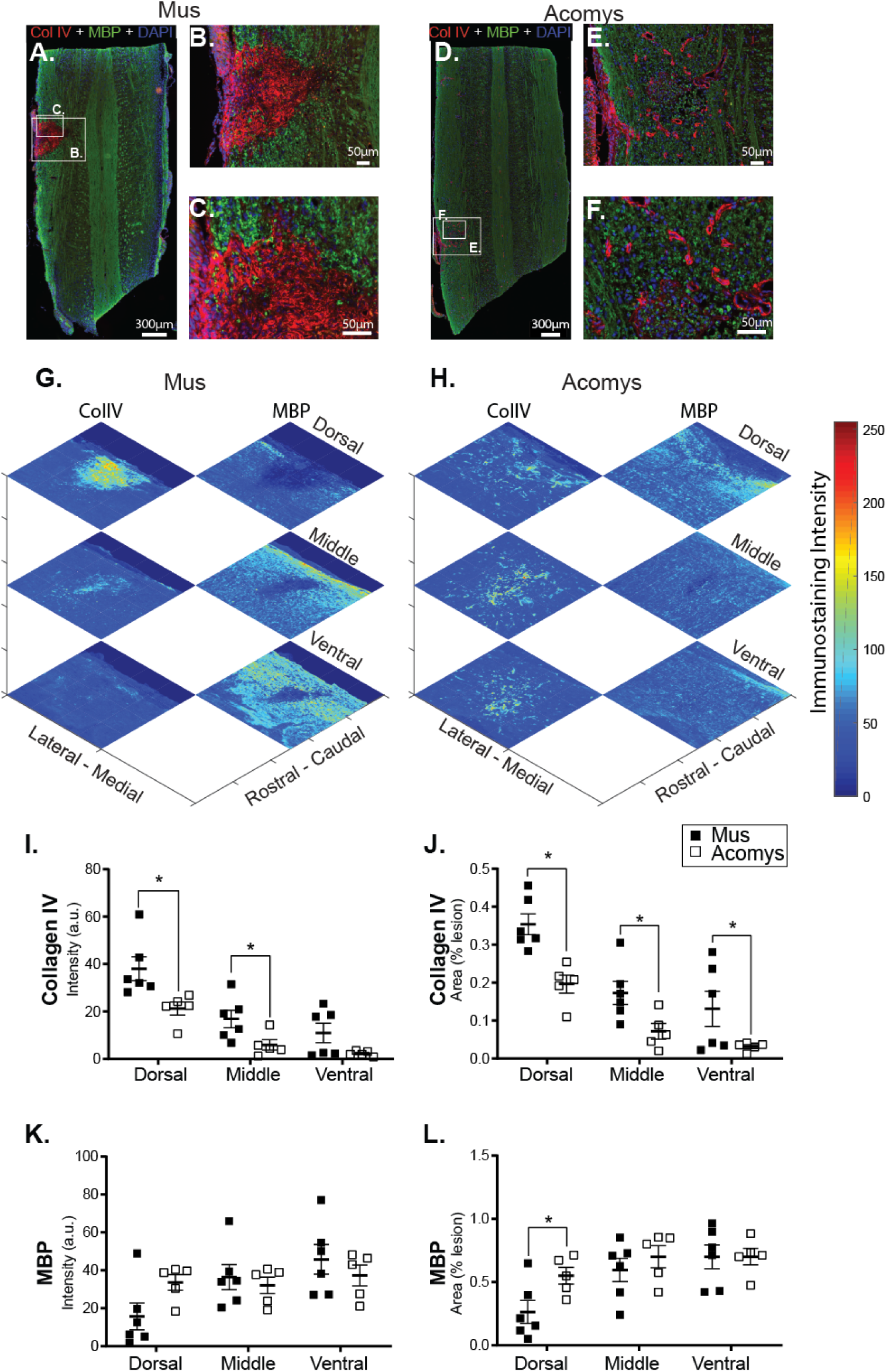
**A-F**. Photomicrographs of collagen IV (red), myelin basic protein (green) and DAPI (blue) staining in the cervical spinal cord (C2-C6) 4 weeks post-SCI in *Mus* (**A-C**) and *Acomys* (**D-F**). **B**,**C**,**E**,**F**. High magnification images of the lesion epicenter shown in panel A for *Mus* (**B**,**C**) and panel D for *Acomys* (**E**,**F**). **G-H**. Representative heat maps of the intensity of collagen IV and myelin staining at the epicenter for each mouse. **I**,**J**. Quantification of the amount of collagen IV staining at the site of the lesion throughout the spinal cord and separated into dorsal, middle, and ventral spinal regions. **K**,**L**. Quantification of the amount of myelin basic protein staining at the site of the lesion throughout the spinal cord and separated into dorsal, middle, and ventral spinal regions. Data shown are means ± standard error. * = p < 0.05.

In contrast to collagen IV, the density of MBP within the spinal lesion appeared greater in *Acomys*. The intensity of MBP staining (a.u.) was similar between species (F(1,27)=0.105, P=0.748), but different across location (F(2,27)=3.52, P=0.0438). Inspection of the data distribution indicated a possible difference in the dorsal spinal cord (Figure 4K). This is more evident when MBP is expressed relative to the lesion area (Figure 4L), as follows: location, F(2,27)=6.644, P=0.005; species, F(1,27)=3.474, P=0.073. Post hoc tests indicate differences in the dorsal spinal cord with greater staining in *Acomys* as compared to *Mus* (P=0.026; Figure 4L).

Photomicrographs of GFAP immunostaining in *Acomys* and *Mus* are provided in Figure 5A-F with heat maps shown in Figure 5G-H. Evaluation of the immunohistochemical sections indicated differences between the two groups. Specifically, in *Mus*, GFAP positive astrocytes had a hypertrophied morphology and were concentrated at the lesion epicenter (Figure 5A), characteristic of reactive gliosis (Wilhelmsson et al., 2004). In *Acomys* GFAP positive astrocytes were organized in a network near the spinal lesion (Figure 5D). The distribution of GFAP staining intensity across all spinal cords is shown in Figure 5I-J. In the middle of the spinal cord, the non-normalized (% of total ipsilesional cord area) GFAP staining intensity suggest a different distribution in *Acomys*. The 2-way ANOVA values for non-normalized data were as follows: species, F(1,30)=2.038, P=0.164; location, F(2,30)=3.251, P=0.053). The 2-way ANOVA values for normalized data were: species, F(1,30)=2.213, P=0.147; location, F(2,30)=2.332, P=0.0115.

Photomicrographs depicting IBA1 immunostaining and associated heat maps are provided in Figure 6A-H. Visual inspection suggested morphological differences between groups. Specifically, in *Mus* spinal cords, IBA1 positive cells demonstrated a large, rounded appearance (Figure 6C) that is consistent with activated macrophages/microglia (Wu et al., 2005). In *Acomys*, IBA1 positive cells were smaller as shown in Figure 6F. The distribution of staining intensity is shown in Figure 6I-J. These data were generally similar between the species, with possible suggestion of reduced IBA1 intensity in the middle spinal cord sections of *Acomys* as compared to *Mus*. The 2-way ANOVA values for non-normalized data were as follows: species, F(1,30)=1.208, P=0.281; location, F(2,30)=0.0362, P=0.965. For the normalized data (% ipsilesional hemicord), the values were: species, F(1,30)=3.17, p=0.085; location, F(2,30)=0.374, P=0.691.

**Figure 5:**
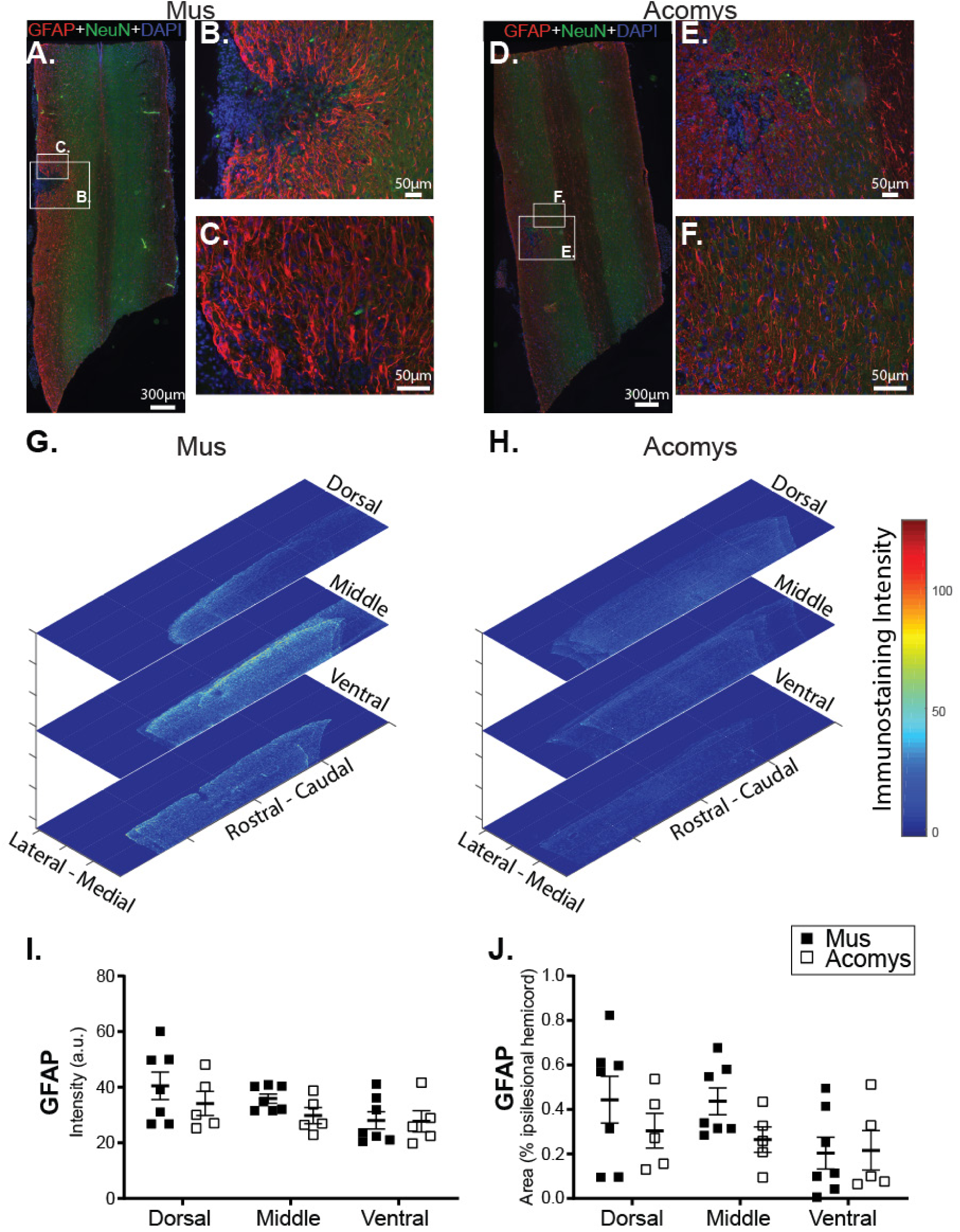
**A-F**. Photomicrographs of GFAP (red), NeuN (green) and DAPI (blue) staining in the cervical spinal cord (C2-C6) 4 weeks post-SCI in *Mus* (**A-C**) and *Acomys* (**D-F**). **B**,**C**,**E**,**F**. High magnification images of the lesion epicenter shown in panel A for *Mus* (**B**,**C**) and panel D for *Acomys* (**E**,**F**). **G-H**. Representative heat maps of the intensity of IBA1 staining at the epicenter for each mouse. Note: the size of heat map images are the ROI that were used for the analysis. **I**,**J**. Quantification of the amount of collagen IV staining at the site of the lesion throughout the spinal cord and separated into dorsal, middle, and ventral spinal regions.

**Figure 6:**
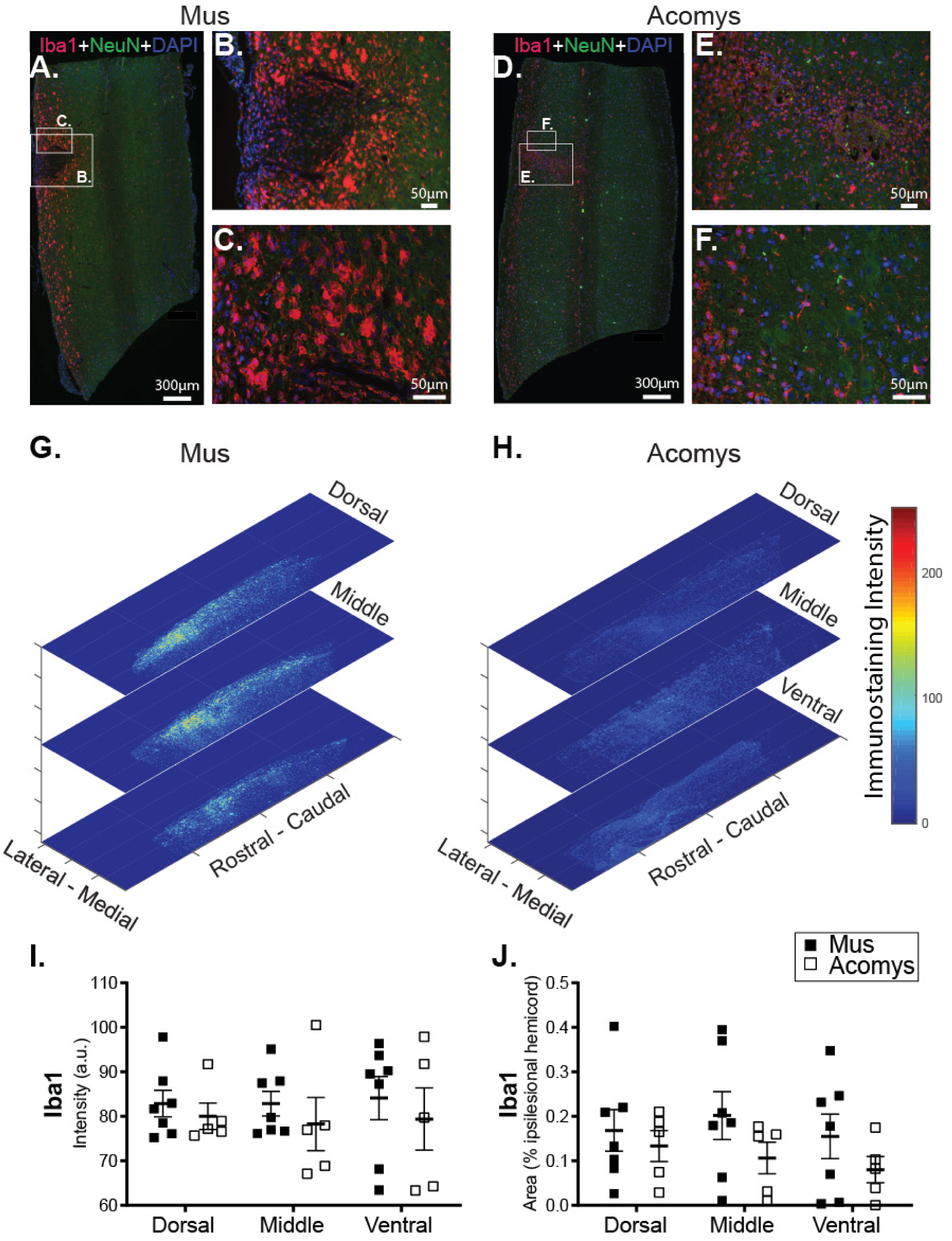
**A-F**. Photomicrographs of IBA1 (red), NeuN (green) and DAPI (blue) staining in the cervical spinal cord (C2-C6) 4 weeks post-SCI in *Mus* (**A-C**) and *Acomys* (**D-F**). **B**,**C**,**E**,**F**. High magnification images of the lesion epicenter shown in panel A for *Mus* (**B**,**C**) and panel D for *Acomys* (**E**,**F**). **G-H**. Representative heat maps of the intensity of IBA1 staining at the epicenter for each mouse. **I**,**J**. Quantification of the amount of collagen IV staining at the site of the lesion throughout the spinal cord and separated into dorsal, middle, and ventral spinal regions.

## DISCUSSION

These experiments indicate that the African spiny mouse, *Acomys*, has a unique molecular and immunohistochemical response to SCI. In specific, three times the number of neurogenesis related genes were induced in *Acomys* as compared to *Mus* and these results stand in sharp contrast to the wound healing arrays where many pro-inflammatory and fibrosis related genes were induced only in *Mus*. The *Acomys* reaction to SCI is consistent with a reduced inflammation and fibrosis observed in our studies on skin regeneration using wound healing arrays (Brant et al., 2015), antibody arrays (Brant et al., 2016) and transcriptome analyses (Brant et al., 2019). Thus, we conclude that the blunted immune response and lack of fibrosis in *Acomys* also extends to the injured spinal cord and may be a feature of any tissue damage inflicted upon this species. In contrast, the *Mus* response is typical of wounding in most mammals, namely the generation of a pro-inflammatory environment by the production of cytokines secreted by leukocytes, neutrophils, macrophages, microglia, and fibroblasts. These cytokines attract additional blood cells and macrophages to the wound site and can assist with the induction of neurotrophins (Fan et al., 2018), but excess levels cause scarring (Werner & Grose, 2003). This self-reinforcing system can induce fibrosis and subsequent inhibition of axonal regrowth across a spinal lesion. Conversely, in the absence of this “cytokine storm”, as it has been referred to (Tisoncik et al., 2012), the spinal environment in *Acomys* may be more conducive to regeneration.

In addition to the induction of pro-inflammatory cytokines, the majority of the remaining genes in *Mus* can be linked together through transforming growth factor-β (TGFβ) and the extracellular matrix (ECM) in the development of a fibrotic response. One of the most highly upregulated genes in *Mus* was *Timp1*, which functions to inhibit the action of the matrix metalloproteases. The same phenomenon was seen in *Mus* skin wound arrays (Brant et al., 2016), suggesting that the fibrotic response of *Mus* to wounding involves the inhibition of MMPs which may generate a more rigid ECM. The interaction of this rigid matrix with the intracellular compartment occurs via the integrins and the upregulation of integrins (e.g. *Itgα5*, along with Itgα1 – a receptor for fibronectin and fibrillin-1), could signal a fibrotic response to cells at the wound site. Activation of the TGFβ signaling pathway is also a key event in fibrosis (Schachtrup et al., 2010). *Serpine1* expression is promoted by TGFβ1 (Honda et al., 2017; Samarakoon & Higgins, 2008) and it is the major physiological regulator of the plasmin based cascade which is involved in fibrotic disorders of the vascular system and several organs such as skin, liver, lung and kidney. *Serpine1* is also itself a pro-inflammatory molecule as it activates macrophages (Gupta et al., 2016). Another plasminogen activator involved in the fibrotic pathway (He et al., 2018) upregulated following SCI in *Mus* was *Plau*, which could supplement the effects of *Serpine1* on fibrosis. In addition to the role these TGFβ1 induced genes play in fibrosis, the activity of plasmin generated by their proteolytic activity is also involved in proteolysis of the ECM which itself could release TGFβ (Deryugina & Quigley, 2012). Interestingly, the *TGF*β*1* gene was induced to the same degree in the spinal tissue of *Mus* and *Acomys* in both the wound and neurogenesis arrays, and also in our skin regeneration experiments (Brant et al., 2015). This observation suggests that post-transcriptional mechanisms are involved in TGFβ1 and fibrotic responses such as its release from the matrix by biomechanical forces or the induction of downstream targets.

In addition to reduced fibrosis-related signaling, following SCI *Acomys* exhibit an upregulation of molecules associated with neurogenesis and regeneration including *Bmp2, Shh, GDNF*, and a tendency for elevated *Notch 1* and *Stat3* levels. This response involves signaling pathways identified in other model systems such as salamanders and zebrafish which successfully regenerate spinal tissue after injury. Specifically, *Wnt, BMP* and *Hh* pathways as well as several growth factors have been implicated in regulating the regenerative capacity of the spinal cord (Vergara, Arsenijevic, & Del Rio-Tsonis, 2005). In mammals, these molecules are known to play important roles during development by establishing signaling gradients which control cellular differentiation, axonal guidance, and neurogenesis (Cardozo et al., 2017).

Consistent with the molecular signature of the acutely injured *Acomys* spinal cord, we found a substantial reduction in staining for collagen IV in the vicinity of the spinal lesion after chronic injury in *Acomys*. Collagen is a major component of the fibrous scar that forms in the injured spinal cord (Hermanns, Klapka, & Müller, 2001; Stichel & Müller, 1998). Collagen can be expressed by multiple cells types, including endothelial cells (Schwab, Beschorner, Nguyen, Meyermann, & Schluesener, 2001), astrocytes (Liesi & Kauppila, 2002) and fibroblasts (Berry et al., 1983), and is generally considered to be part of the barrier to growth of axonal projections after SCI (Klapka & Müller, 2006). Another contribution to the glial scar comes from reactive astrocytes that express GFAP (Bovolenta et al., 1992). Our evaluation of GFAP intensity in the chronically injured *Acomys* spinal cord did not demonstrate a clear difference as compared to *Mus*, although a tendency for reduced staining can be seen in the data. Qualitative differences in the pattern and morphology of GFAP staining around the lesion epicenter are consistent with robust gliosis in *Mus*, which was not as pronounced in *Acomys*. Microglia are also activated after SCI and can be found in increased numbers near the interface of astrocytes and infiltrating leukocytes (Bellver-Landete et al., 2019; Hawthorne & Popovich, 2011). Recent evidence indicates that this microglial/macrophage response may in fact be beneficial to the injured spinal cord (Bellver-Landete et al., 2019; Gensel & Zhang, 2015). In our study, there was no suggestion of a difference in IBA1 staining intensity between the chronically injured *Acomys* and *Mus* spinal cords, which labels both macrophages and microglia (Imai, Ibata, Ito, Ohsawa, & Kohsaka, 1996). However, visual comparison of the IBA1 positive cells reveals *Acomys* have fewer, large, round IBA1 positive cells consistent with phagocytic morphology characteristic of activated microglia/macrophages (Wu et al., 2005).

In summary, the molecular and histologic data presented here are consistent with a growing literature that indicates that *Acomys* has a tissue response to injury that may be relatively unique among mammals. Prior work definitively establishes that *Acomys* does not undergo fibrosis and has remarkable regenerative capacity in dermis, smooth muscle, skeletal muscle, and other tissue following injury (Brant et al., 2019; Brant et al., 2016; Jiang, Harn, Ou, Lei, & Chuong, 2019; Seifert et al., 2012). Our current data support the hypothesis that this unique response to injury extends to the spinal cord. Based on the reduced inflammatory and fibrotic response and indication of enhanced regenerative capacity, we suggest that *Acomys* merits further study as comparative model to study adaptive responses to SCI.

## Acknowledgements

This work was supported by funding from the National Institute of Health, grant numbers: K99 HL143207-01 (KS), 1F32NS095620-01 (KS), 1R01NS080180-01A1 (DF), T32-ND043730 (MS) and F31HL145831 (MS), The Keck Foundation (MM) and R21OD023210 (MM and JOB). We thank Dr. Lila Wollman and Shreya Patel for their assistance with surgical procedures. We also thank Marda Jorgensen for excellent histological and immunochemical work, and Ethan Benevides for imaging work. The authors declare no competing financial interests

