## Supplemental Tables 1-2 for "Molecular and histologic outcomes following spinal cord injury in spiny mice, *Acomys cahirinus*"

**Supplementary Table 1. RT-qPCR oligo sequences.**

***Acomys* RT Primers**

| Gene | Forward Primer Sequence | Reverse Primer Sequence |
|---|---|---|
| ***Serpine*** | AGCGGCCTGCTATGAGATTA | CCTGGTCAACCACCTCTGTT |
| ***Plau*** | CCGTTCCCTCACACCATTAG | TTCATGAGGTCTGCTGTTGG |
| ***Timp1*** | ACGTCCAGCGATGAGAAACT | GTTCGAAATTGTGGGGAATG |
| ***Itgb5*** | TGGCACTGAACAGTGAGGAC | AGCCAGGTGTACTGCAAGGT |
| ***Gapdh*** | CGACCTTCACCATCTTGTCA | CCCACCAACCTGGTTCCTAT |
| ***Bmp2*** | TCTTTTCCATTCCATCCCATA | AAACATCCCTGCTCCCTTCT |
| ***Gdnf*** | AAGCGCTGCCACTTGTTTAT | TGACCAGCGACTCCAATATG |
| ***Shh*** | TTTACACTTGCGGAAAAGCA | ATTTTGTGAGGCCAAGCAAC |
| ***Stat3*** | AAAAGTGCAGTGCCAGGAGT | TCAGCAGCTCCAGGGTTATT |
| ***Tgfb1*** | GAGCGCACGATCATGTTG | CTGCCCCTACATCTGGAGTC |

***Mus* RT Primers**

| *Gene* | Forward Primer Sequence | Reverse Primer Sequence |
|---|---|---|
| ***Serpine*** | CACGGTACCTTCTTGTGCAG | GGGGATGAAAGAGACAGCAG |
| ***Plau*** | TGAGATCTGCTGTTGGGAAA | TCCCTCAAGCCGTTAGTGTC |
| ***Timp1*** | CTTGGTTCCCTGGCGTACTC | ACCTGATCCGTCCACAAACA |
| ***Itgb5*** | GGCCAGCTCCGTCTATGATG | TGGATGTCTGAGCCATTAAGGA |
| ***Gapdh*** | TGACCTCAACTACATGGTCTACA | CTTCCCATTCTCGGCCTTG |
| ***Bmp2*** | TCTTGTGGGCCCTCATAAAG | GCGTGTCAGGCAAATGTAGA |
| ***Gdnf*** | TCCTGACCAGTTTGATGACG | CCTGCCGATTCCTCTCTCTT |
| ***Shh*** | GTGGCGGTTACAAAGCAAAT | CATGGGGGTCCACAAATTAT |
| ***Stat3*** | TTCAGACGATATGGGGTTCG | ATGGATCTGACCTCGGAGTG |
| ***Tgfb1*** | TGCCCTCTACAACCAACACA | GTTGGACAACTGCTCCACCT |

**Supplemental Table 2: A.** Differentially expressed genes in the ipsilateral spinal cord (C2-C6) 3 days post-SCI compared to spinal intact controls within each species using a wound healing array. Only genes with a fold change >1.5 are recorded. **B.** Differentially expressed genes in the ipsilateral spinal cord (C2-C6) 3 days post-SCI compared to spinal intact controls within each species using a neurogenesis array. Only genes with a fold change >1.5 are recorded.

A. Wound Healing Array

| | Mus | Acomys |
|---|---|---|
| Serpine1 | 37.05 | |
| Timp1 | 28.16 | -4 |
| Il6 | 9.88 | |
| Cxcl3 | 8.57 | -2.28 |
| Ccl12 | 6.02 | |
| Ccl7 | 5.4 | |
| Tnf | 4.52 | 2.45 |
| Itga5 | 4.25 | |
| Plaur | 4.24 | |
| Il1b | 3.94 | |
| Mmp1a | 3.57 | |
| Tagln | 2.83 | -4.94 |
| Col5a3 | 2.3 | |
| Plau | 2.18 | |
| Tgfb1 | 1.84 | |
| Hgf | 1.69 | |
| Mapk3 | -1.51 | |
| Itga4 | -1.52 | |
| Rhoa | -1.52 | -1.56 |
| Il6st | -1.54 | -189.27 |
| Wnt5a | -1.75 | |
| Tgfa | -1.77 | |
| Tgfbr3 | -1.84 | |
| Actc1 | -1.96 | |
| Itga1 | -1.98 | |
| Mif | -2.02 | -1.79 |
| Pdgfa | -2.09 | |
| Egf | -2.14 | |
| Pten | -2.14 | -1.97 |
| Mmp9 | -2.17 | |
| Mapk1 | -2.25 | -2.24 |
| Vtn | -2.59 | -160.44 |
| Il4 | -2.6 | |
| Vegfa | -2.68 | -1.93 |
| Plg | -2.78 | |
| Fgf10 | -3.27 | |
| Col4a3 | -3.42 | -3.02 |
| Col5a2 | | 14.58 |
| Cdh1 | | 3.34 |
| Ctgf | | -2.11 |
| Il2 | | -2.14 |
| Itgb6 | | -2.31 |
| Col14a1 | | -2.82 |
| Itgb5 | | -3.41 |
| Angpt1 | | -3.59 |
| F13a1 | | -6.27 |
| Ptgs2 | | -6.36 |
| Rac1 | | -13.82 |
| Ctsl | | -16.64 |
| Ctnnb1 | | -18.93 |
| F3 | | -28.47 |
| Itgav | | -36.73 |

B. Neurogenesis Array

| | Mus | Acomys |
|---|---|---|
| S100a6 | 6.96 | |
| Tgfb1 | 3.43 | 3.6 |
| Flna | 2.98 | 5.5 |
| Gdnf | 2.55 | 3.57 |
| Fgf2 | 2.15 | |
| Erbb2 | 1.75 | |
| Stat3 | 1.54 | 2.93 |
| Mdk | | -2.47 |
| Bcl2 | | 1.7 |
| Sox2 | | 2.08 |
| Ptn | | 1.68 |
| Creb1 | | 1.58 |
| Tnr | -1.51 | |
| Adora2a | -1.52 | |
| Hdac4 | -1.52 | |
| Shh | -1.53 | 1.73 |
| Dvl3 | -1.53 | 1.5 |
| Hey1 | -1.54 | |
| Olig2 | -1.54 | |
| Pard3 | -1.55 | |
| Adora1 | -1.55 | |
| Artn | -1.56 | |
| Bmp2 | -1.58 | 2.99 |
| Nf1 | -1.58 | |
| App | -1.61 | |
| Egf | -1.64 | |
| Pax5 | -1.66 | |
| Ndn | -1.68 | |
| Pou3f3 | -1.75 | |
| Apbb1 | -1.81 | |
| Pafah1b1 | -1.84 | -1.68 |
| Dcx | -1.84 | 1.51 |
| Robo1 | -1.87 | 1.61 |
| Map2 | -1.88 | |
| Tenm1 | -1.9 | |
| Slit2 | -1.91 | |
| Dlg4 | -1.93 | |
| S100b | -1.98 | |
| Ache | -2.02 | |
| Vegfa | -2.03 | |
| Grin1 | -2.04 | |
| Dll1 | -2.04 | |
| Nrg1 | -2.07 | |
| Cdk5r1 | -2.08 | |
| Drd2 | -2.08 | |
| Neurod1 | -2.08 | |
| Heyl | -2.4 | 1.86 |
| Chrm2 | -2.54 | |
| Neurog2 | -3.8 | |
| Efnb1 | | 2.35 |
| Ntn1 | | 1.93 |
| Notch1 | | 1.87 |
| Notch2 | | 1.77 |
| Ascl1 | | 1.72 |
| Ndp | | 1.62 |
| Mef2c | | 1.58 |
| Bdnf | | 1.57 |
